# Dectin-2-dependent adaptive immunity governs intestinal clearance of systemic *Candida albicans*

**DOI:** 10.64898/2026.01.28.702262

**Authors:** Ivy M. Dambuza, Fabián Salazar, Jamie Harvey, Annie Phillips-Brooks, Daniel H. Kaplan, Gordon D. Brown

## Abstract

Effective immunity to *Candida albicans* requires coordination between innate recognition and induction of adaptive CD4 T cell responses. While the C-type lectin receptor Clec4n (Dectin-2) is known to drive Th17 polarization, its role in shaping tissue-specific adaptive responses remains incompletely understood. Here, we used an OT-II antigen-specific CD4 T cell transfer model combined with OVA-expressing *Candida albicans* to dissect the function of Dectin-2 during systemic infection. We found that Dectin-2 is dispensable for antigen presentation and CD4 T cell priming in gut-draining lymph nodes. Moreover, we show that Dectin-2-deficient mice fail to control fungal growth in the intestinal mucosa, despite elevated local production of IL-17A and GM-CSF. The increased susceptibility of the Dectin-2-deficient mice was associated with impaired neutrophil activation in the intestinal mucosa. These findings identify a tissue-specific checkpoint role for Dectin-2, linking balanced adaptive Th17 cytokine responses to granulocyte function, and revealing a previously unappreciated mechanism required for anti-fungal immune regulation at intestinal mucosal surface.

## Introduction

Systemic candidiasis caused by *Candida albicans* accounts for up to 40% of fungal sepsis cases and carries case-fatality rates exceeding 45% despite anti-fungal therapy (1). Effective clearance from mucosal sites depends not only on robust innate immunity but crucially on adaptive CD4 T cell responses (2, 3). Among the pattern recognition receptors (PRRs) that bridge these immune arms, Clec4n (Dectin-2) has emerged as a key C-type lectin receptor (CLR), selectively recognizing α-mannans on the *C. albicans* cell wall (4, 5). Dectin-2 is expressed pre-dominantly on dendritic cells (DCs) and macrophages, and signals through the FcRγ-Syk-CARD9-MALT1 axis upon ligand engagement (6). This signalling cascade leads to the induction of pro-inflammatory cytokines including IL-1β, IL-6, and IL-23, which are essential for the polarization of naïve CD4 T cells toward a Th17 fate (7). Th17 responses, characterized by cytokines such as IL-17A production, play a pivotal role in neutrophil activation and recruitment, resulting in clearance of *Candida* from infected tissues (14–17).

Several studies have demonstrated that mice lacking Dectin-2 or those treated with Dectin-2-blocking antibodies exhibit impaired Th17 differentiation and reduced IL-17 production during *C. albicans* infection, resulting in compromised fungal clearance and increased susceptibility (5, 8). Notably, while Dectin-2 predominantly drives Th17 responses, it also contributes to Th1 immunity in coordination with other CLRs such as Clec7a (Dectin-1) (7). Mechanistically, Dectin-2 activates selective c-Rel-dependent transcriptional programs in DCs that shape anti-fungal Th17 responses (9). Others have shown that Dectin-2 cooperates with Clec4e (Mincle) to fine-tune anti-fungal T helper polarization (10). Additionally, Dectin-2 signalling has been linked to “trained immunity,” where innate cells exposed to fungal mannans develop heightened responsiveness and potentiate subsequent adaptive responses (11).

Despite the substantial evidence linking Dectin-2 to the generation of anti-fungal T cell responses, the precise role of Dectin-2 in regulating antigen-specific CD4 T cell function, particularly within the gastrointestinal tract, remains poorly defined. Most studies have relied on bulk cytokine measurements or *ex vivo* non-specific restimulation of spleen polyclonal T cells, which do not fully capture the dynamics of bona fide antigen-specific responses at mucosal sites. As a result, it remains unclear whether Dectin-2 is required during the priming, effector, or recall phases of adaptive immunity, particularly within tissue compartments that serve as reservoirs for fungal dissemination, such as the gastrointestinal tract (GIT). We previously showed that impaired fungal control in the GIT of Dectin-2 knockout (KO) mice was not associated with antigen-specific T cell loss, unlike other CLRs such as Dectin-1 (12), suggesting a distinct immunoregulatory mechanism downstream of Dectin-2. In this study, we demonstrate that Dectin-2 is critical for the optimal activation of granulocytes in the GIT during systemic candidiasis and uncover a non-redundant role for Dectin-2 in bridging a balanced IL-17A and GM-CSF response with innate effector functions. This underscores the significance of Dectin-2 as a tissue-specific checkpoint in systemic fungal infections.

## Results

### Dectin-2 deficiency impairs intestinal fungal clearance in the presence of antigen specific CD4 T cells

Our previous work demonstrated that Dectin-2 deficiency does not result in loss of antigen-specific CD4 T cells during *C. albicans* infection (12). However, whether Dectin-2 is required for CD4 T cell priming, effector function, or recall responses in the GIT remains unresolved. Since the GIT is a key anatomical reservoir for *C. albicans* colonization and dissemination during systemic candidiasis, understanding the specific contribution of Dectin-2 to adaptive immune control in this compartment is critical. To address this, we used our previously employed model (12) involving adoptive transfer model of antigen-specific CD4 T cell immunity, in which naïve OT-II cells are introduced in wild-type (WT) and Dectin-2 KO mice, followed by systemic infection with an OVA-expressing *C. albicans* strain (Calb-Ag) (13). This allowed us to directly evaluate the impact of Dectin-2 on antigen-specific T cell responses during systemic fungal infection, with particular focus on the GIT.

We first assessed whether the innate functions of Dectin-2 were necessary for fungal control, in the absence of antigen-specific CD4 T cells. In mice that did not receive OT-II T cell transfer, fungal burdens in the intestines were comparable between Dectin-2 KO mice and WT mice at both day 3 and day 6 post-infection (**Figures 1A** and **B**). This suggests that under baseline innate immune conditions, Dectin-2 is not essential for intestinal fungal clearance, indicating some redundancy in pattern recognition receptor (PRR) pathways during early fungal containment. However, the functional requirement for Dectin-2 became strikingly apparent in the presence of antigen-specific T cells. When OT-II T cells were transferred prior to infection, WT mice were able to completely clear intestinal fungal burdens by day 6 (**Figures 1C** and **D**). In sharp contrast, Dectin-2 KO mice failed to eliminate fungal burden from the GIT; instead, fungal burdens persisted and had even increased by day 6 (**Figures 1C** and **D**). Fungal burdens in the kidney were comparable between Dectin-2 KO and WT mice at both day 3 and day 6, regardless of OT-II T cell transfer (**Figures 1A-D**), consistent with our previous observations that antigen-specific CD4+ cells are not recruited to the kidney during systemic candidiasis (25). These data indicate a failure of adaptive immune-mediated fungal control in the absence of Dectin-2 specifically in the GIT.

**Figure 1.**
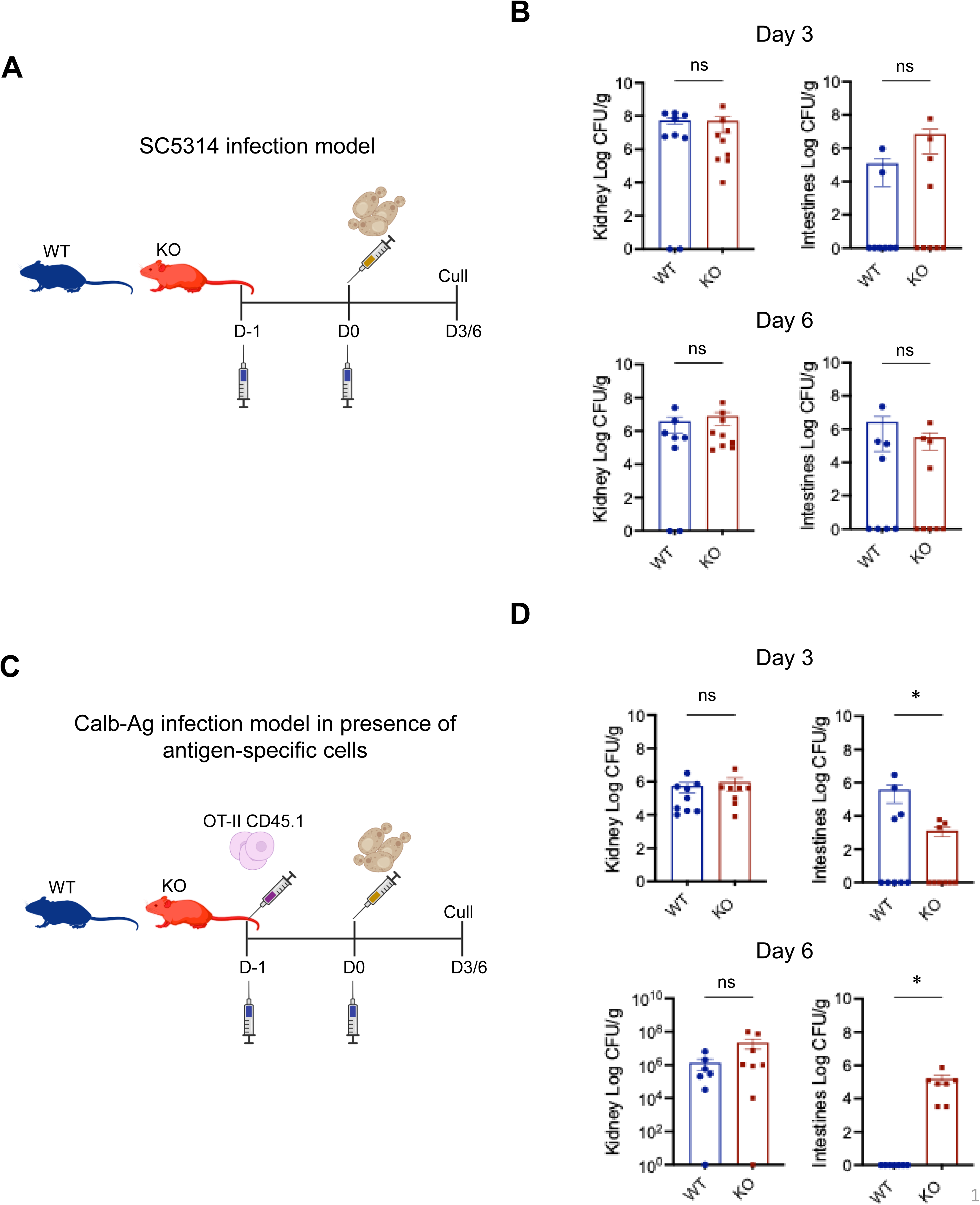
Dectin-2 deficiency impairs intestinal fungal clearance in the presence of antigen specific CD4 T cells. Fungal burdens in the kidneys or intestine at day 3 or 6, as indicated, in wild-type (WT) and Dectin-2 KO mice were challenged systemically with OVA-expressing *C. albicans* (Calb-Ag) in the absence **(A,B)** or presence **(C,D)** of adoptively transferred OT-II CD4 T cells. Data represent mean ± SEM from n = 4-5 mice per group (pooled 2 experiments); statistical significance was determined by *t*-test, p < 0.05 is (*) considered significant, ns = not significant.

### Dectin-2-deficient dendritic cells show intact co-stimulatory molecule expression in gut-draining lymphoid tissue

Given the impaired intestinal fungal clearance observed in Dectin-2 KO mice in the presence of antigen-specific CD4 T cells (**Figure 1**), we sought to investigate whether this defect could reflect altered DC activation or antigen presentation capacity. DCs are central to initiating adaptive immunity, providing both MHC-peptide complexes and co-stimulatory signals essential for naïve CD4 T cell priming. Dectin-2 has been shown to regulate DC maturation, cytokine production, and expression of co-stimulatory molecules following fungal stimulation (7). However, Dectin-2’s contribution to DC function during systemic infection has not been clearly defined, especially in the gut-draining mesenteric lymph nodes (mLN). We analyzed the expression of CD40, CD80, and CD86 on CD11c+MHC-II+ DCs in the mLN at days 3 and 6 post-infection with Calb-Ag. These timepoints were selected to capture the priming and early activation of CD4 T cell responses. At days 3 and 6, no significant differences were observed overall. The only change detected was a modest reduction at day 6 in the number of CD11c⁺MHC-II⁺ DCs expressing CD40 in the mLNs of Dectin-2 KO mice (**Figure 2A-B**). Similarly, across both timepoints, the mean fluorescence intensities (MFIs), reflecting the level of cellular expression, of all co-stimulatory markers remained unaltered (**Figure 2C-D**), suggesting that Dectin-2 does not regulate these molecules at least in mLN CD11c+MHC-II+ DCs. There was also no alteration in expression of MHCII on these cells (**Figure 2C-D)**

**Figure 2.**
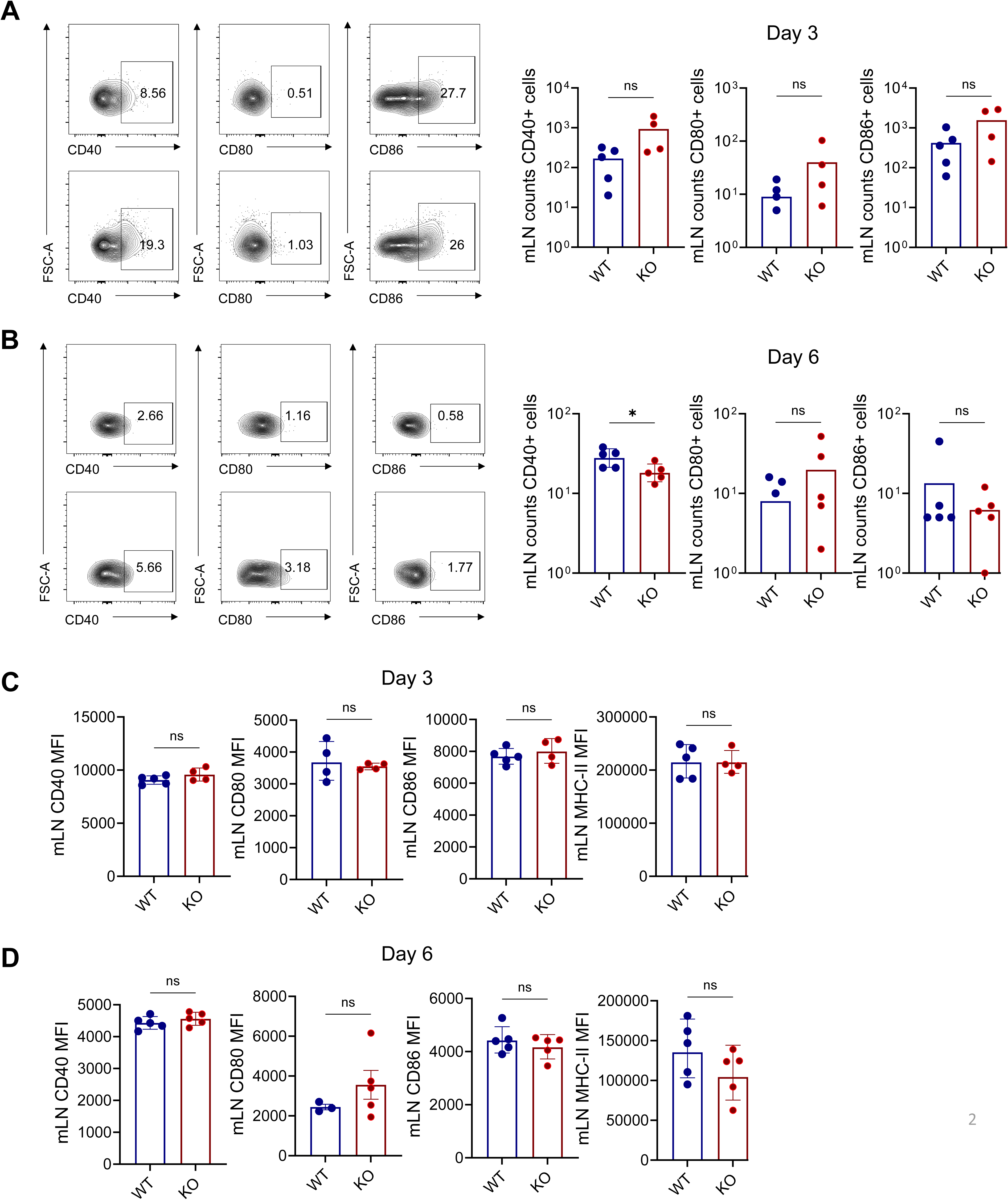
Dectin-2-deficient dendritic cells show intact co-stimulatory molecule expression in gut-draining lymphoid tissue. Surface expression and counts of CD80, CD86, CD40, and MHC-II on dendritic cells (DCs) from intestinal draining lymphoid tissue (mLN) were analyzed by flow cytometry at days 3 and 6 post-infection. **(A)** Day 3, **(B)** day 6, co-signalling molecules **(C,D)** Mean fluorescence intensity (MFI) values for these markers. **(Supplementary Fig 2A-D)** Surface expression and counts of CD80, CD86, CD40, and MHC-II on DCs from splenic DCs were analyzed by flow cytometry at days 3 and 6 post-infection. **(Supplementary Figure 2E)** Gating strategy DC characterisations. Data represent mean ± SEM, n = 4-5 per group; *t*-test, p < 0.05 is (*) considered significant, ns = not significant.

To determine whether this phenotype was restricted to the mLN or reflected a systemic response, we also analyzed DCs in the spleen. At day 3 and day 6, we observed no major changes in the counts of DCs expressing CD40, CD80 and CD86 in the absence of Dectin-2 (**Supplementary Figure 2A, B**). There were also no significant changes in the expression levels of these molecules (MFI), except for MHC-II where there was a consistent lower level of expression in splenic DCs from Dectin-2 KO mice (**Supplementary Figure 2C, D, E**). These findings suggest that T cell priming signals are largely preserved on DC in the absence of Dectin-2.

### Dectin-2 is not required for antigen-specific CD4 T cell priming or Th1/Th17 polarization in gut-draining lymph nodes or lamina propria

Given that Dectin-2 has been shown to support Th17 immunity (3), we next investigated whether the impaired intestinal fungal clearance observed in Dectin-2 mice could be attributed to defects in antigen-specific CD4 T cell responses. In our previous study we showed that lack of Dectin-2 did not impact survival or proliferation of antigen-specific CD4 T cells, at least at day 3 post infection (12). Here we assessed the expansion and activation of OT-II cells in the mLN and lamina propria at day 6 after *C. albicans* infection. Similar to what we had observed previously at day 3 (12), the frequency and activation (shown by CD44+ and CD69+) of OT-II cells in the mLN was similar between both genotypes at day 6 (**Figure 3A,B and Supplementary Fig3**). Because Dectin-2 has been strongly linked to development Th17 and Th1 anti-fungal responses, we then examined whether CD4 T cell differentiation into Th1 (Tbet) or Th17 (RORγt) lineages was altered at day 6. We found no differences in the frequencies of antigen-specific cells expressing RORγt, Tbet and Foxp3 at this later time point (**Figure 3C**), consistent with our previous study (12). Overall, these results indicate that Dectin-2 is not essential for T cell activation of Th17 or Th1 lineage commitment in gut-draining lymphoid tissue.

**Figure 3.**
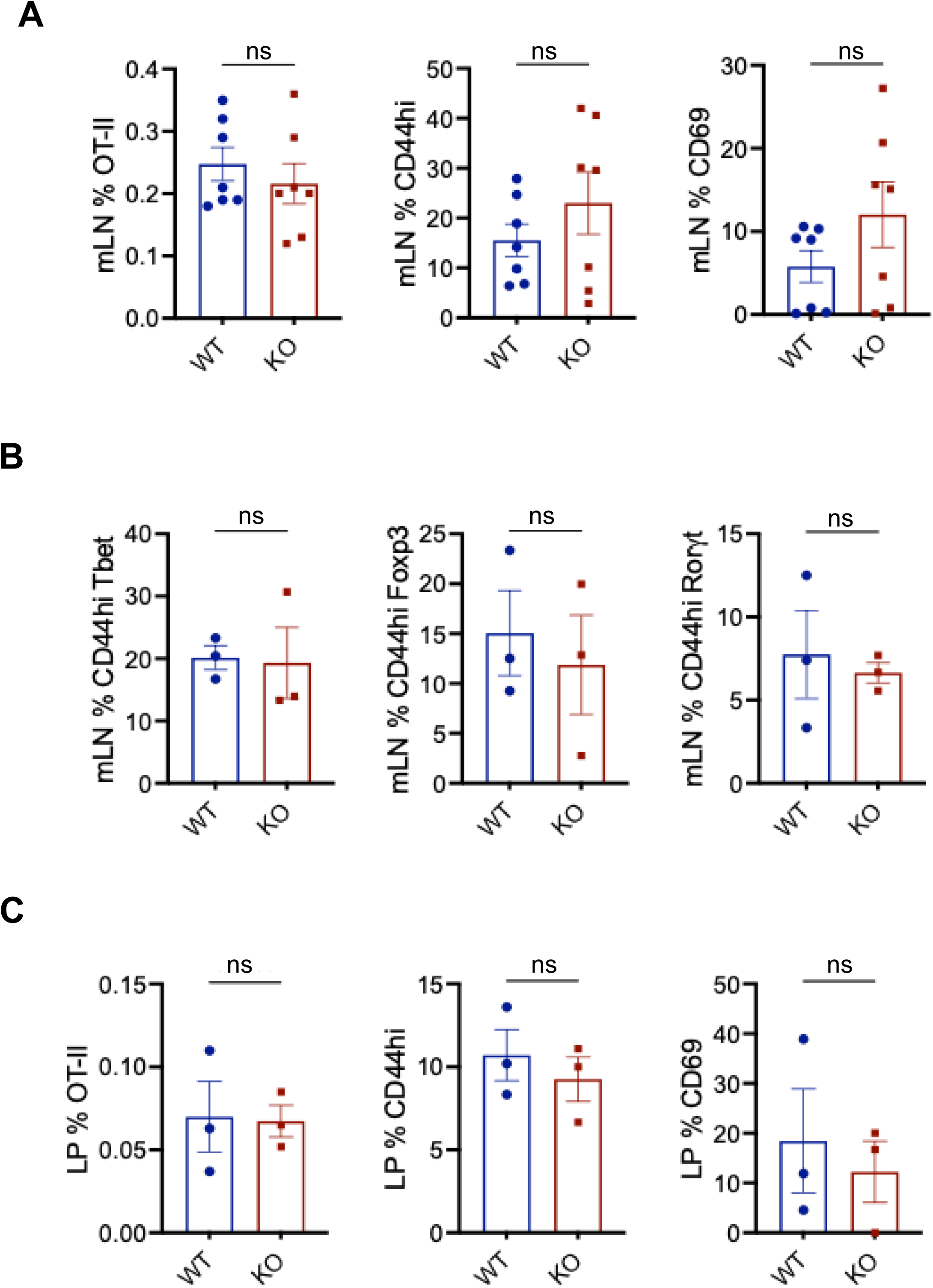
Dectin-2 is not required for antigen-specific CD4 T cell priming or Th1/Th17 polarization in gut-draining lymph nodes or lamina propria. OT-II CD4 T cell activation and expression of Th1 (Tbet) and Th17 (RORγt) transcription factors were assessed by flow cytometry in mesenteric mLN and lamina propria at day 6 post-infection between WT and Dectin-2 KO mice. **(A)** The percentage and activation status (CD44, CD69) of OT-II cells in mLN. **(B)** Frequencies of Tbet+, Foxp3+, and RoRγt+ OT-II cells in the mLN. **(C)** Frequencies of lamina propria (LP) % OTII, CD44+ and CD69+ cells. **Supplementary Figure 3**. Gating strategy to isolate out OTII+ cells and OTII cells expressing CD44 and CD69. Data are shown as mean ± SD, n = 3-5 per group; *t*-test, p < 0.05 is (*) considered significant, ns = not significant.

To evaluate whether Dectin-2 might affect CD4 T cell recruitment at the site of infection, we assessed the frequency of OT-II cells in the intestinal lamina propria (LP). The very low frequency of OT-II cells in the LP prevents polarization analyses of these cells. Nonetheless, we noted comparable recruitment to the LP, and similar activation (**Figure 3C, Supplementary Fig3**), suggesting that effector T cells could be successfully mobilized to the site of infection even in the absence of Dectin-2. Together, these findings demonstrate that the absence of Dectin-2 does not impair T cell expansion or differentiation in secondary lymphoid tissues or intestinal effector sites, pointing to downstream defects in anti-fungal effector function rather than a failure in early adaptive programming.

### Dectin-2 KO mice show elevated IL-17A and GM-CSF levels in the intestine despite impaired fungal clearance

We wanted to determine if the failure to control fungal growth in the intestine stemmed from defects in effector cytokine production, particularly those associated with Th17 responses. IL-17A and GM-CSF are hallmark effector cytokines produced by Th17 cells that coordinate fungal immunity through neutrophil recruitment, granulopoiesis, and activation of tissue-resident phagocytes (14–17). Dectin-2-Syk-CARD9 signalling has previously been shown to be essential for initiating Th17 polarization and IL-17 production following *C. albicans* infections (7–9). Importantly, GM-CSF also enhances anti-fungal defences by licensing monocyte-derived cells and neutrophils to generate reactive oxygen species (ROS) and other anti-microbial functions (18). We therefore quantified these two cytokines and other key pro-inflammatory mediators in intestinal tissue homogenates at day 6 post-infection. Unexpectedly, we observed significantly elevated levels of GM-CSF and IL-17A and increased in Dectin-2 KO mice compared to WT mice (**Figure 4**). Other cytokines, including IFN-γ, IL-4, IL-5, and IL-6, showed no significant differences between the groups. In contrast, IL-1β levels were significantly reduced in Dectin-2 KO mice (**Figure 4**). Together, these results indicate a selective rather than global alteration in cytokine production within the intestinal tissue during systemic candidiasis. Specifically, Dectin-2 appears to be required to restrain IL-17A and GM-CSF responses, while also supporting optimal IL-1β production.

**Figure 4.**
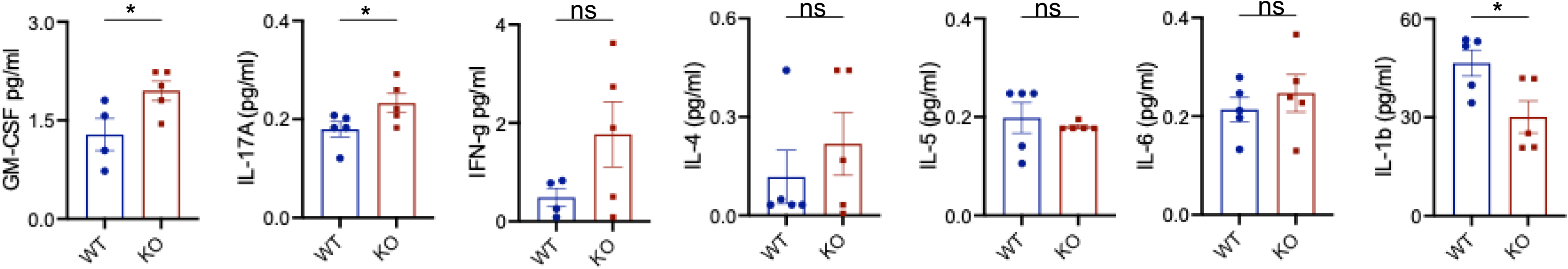
Dectin-2 KO mice show elevated IL-17A and GM-CSF levels in the intestine despite impaired fungal clearance. Cytokine concentrations in intestinal homogenates were measured at day 6 post-infection by ELISA or multiplex assay. Data are mean ± SD n = 5 per group; *t-*test, p < 0.05 is (*) considered significant, ns = not significant.

### Dectin-2 KO mice exhibit increased granulocyte recruitment but impaired neutrophil activation in the intestine

Given the elevated levels of Th17-associated cytokines, IL-17A and GM-CSF, and reduced IL-1β at day 6 post-infection (**Figure 4**), we then examined whether granulocyte dysfunction could explain the failure of Dectin-2 KO mice to effectively control fungal growth in the GIT. Previous studies have implicated Dectin-2 signalling in neutrophil activation (19) and infiltration into tissues (20), linked with effective anti-fungal immunity. These responses often involve regulation in integrin expression or degranulation in similar pathways, although direct evidence for upregulation of CD11b/CD18 and CD63 in GIT following *Candida* infection downstream of Dectin-2 remains to be established. To assess this, we quantified the numbers neutrophils, eosinophils, monocytes, and macrophages in the GIT lamina propria (LP) at day 3 and 6 post-infection and analysed their activation state based on expression of CD18, CD11b and CD63. The frequencies of neutrophils, monocytes and macrophages were not significantly altered at either time point (**Figures 5A-B** and **Supplementary Figure 5A**). In contrast, we observed a slight but reproducible significant increase in eosinophils in mice lacking Dectin-2 compared to WT mice at day 6 post-infection (**Figures 5A-B**), suggesting enhanced recruitment of these phagocytes. Despite robust infiltration, at both time points neutrophils in Dectin-2 KO mice displayed significantly reduced expression of CD18 (integrin β2), a key adhesion molecule involved in trans-endothelial migration and immune synapse formation with fungi (21–23) (**Figure 5C-D**). This indicates impaired neutrophil activation, consistent with functional hypo-responsiveness. Conversely, eosinophils from Dectin-2 KO mice showed increased expression of marker of degranulation, CD63, at day 6 when compared to WT mice (**Figure 5D**). These data suggest that in the absence of Dectin-2, the activation of eosinophils becomes dysregulated. Overall, these results show that Dectin-2 deficiency leads to increased intestinal recruitment of eosinophils without changes in neutrophils, monocytes or macrophages, accompanied by reduced CD18 expression on neutrophils and elevated CD63 expression on eosinophils

**Figure 5.**
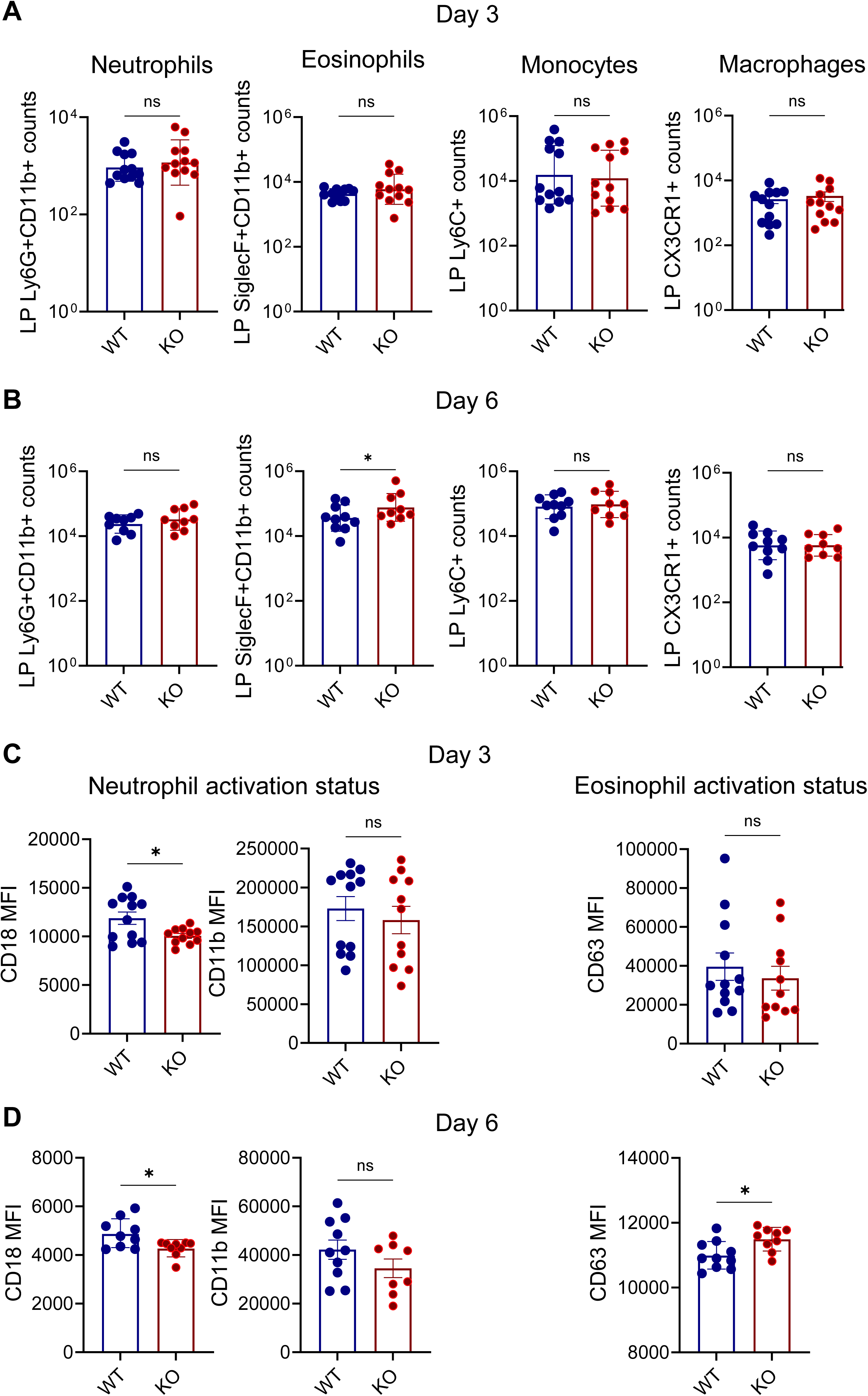
Dectin-2 KO mice exhibit increased granulocyte recruitment but impaired neutrophil activation in the intestine. Flow cytometric quantification of neutrophils, eosinophils, monocytes, and macrophages in intestinal tissue at days 3 and 6 post-infection. **(A,B)** Cellular counts in lamina propria at day 3 and 6 post-infection in Dectin-2 KO mice. **(C,D)** Neutrophil and eosinophil activation in lamina propria at day 3 and 6 post-infection in Dectin-2 KO mice **(Supplementary Figure 5A)** Gating strategy immunophenotyping. Data represent mean ± SEM, n = 4-5 per group (pooled 2 experiments); *t*-test, p < 0.05 is (*) considered significant, ns = not significant.

## Discussion

This study identifies a previously underappreciated tissue-specific role for Dectin-2 in coordinating adaptive anti-fungal immunity within the intestinal mucosa. Although earlier work has established Dectin-2 as a key C-type lectin receptor for driving Th17 responses and promoting fungal clearance during the later stages of systemic *C. albicans* infection (5, 7, 24), its phase-specific contributions to CD4 T cell-mediated anti-fungal defence, particularly in the GIT, have remained poorly defined. Here, we used a targeted approach involving adoptive transfer of antigen-specific OT-II CD4 T cells and infection with an ovalbumin-expressing *C. albicans* strain to interrogate how Dectin-2 influences adaptive responses at tissue sites during fungal dissemination.

Our findings show that defects in fungal clearance in Dectin-2-deficient mice emerged only in the presence of antigen-specific T cells, indicating that Dectin-2 is not required for innate control in the GIT. This finding is consistent with previous studies reporting reduced survival in Dectin-2–deficient mice during long-term infections following exposure to a sublethal infectious dose (8). Despite effective recruitment and polarization of OT-II CD4 T cells toward both Th1 and Th17 fates in Dectin-2 deficient mice and preserved upregulation of co-stimulatory and MHC-II molecules on dendritic cells in gut-draining lymph nodes, fungal clearance from the intestinal tissue was impaired. Together, these findings suggest that while Dectin-2 is not required for the initial priming of CD4 T cells it may play an important role in shaping the quality of the effector response. The elevated IL-17A and GM-CSF levels observed in intestinal homogenates of Dectin-2 deficient mice point to a regulatory function of Dectin-2 in restraining or calibrating T cell cytokine output during anti-fungal immunity. These cytokines are classically associated with Th17 effector function and are known to mobilize and activate granulocytes, especially neutrophils. We found that both immune cell recruitment to the lamina propria and the activation states of neutrophils and eosinophils were altered in the absence of Dectin-2. While neutrophil recruitment to the lamina propria was unchanged, eosinophil infiltration was slightly increased in Dectin-2–deficient mice. In addition, neutrophils displayed impaired activation, whereas eosinophils showed reduced activation at later time points. The contribution of eosinophils to intestinal anti-fungal immunity remains poorly defined, but our data highlight the need for future studies to delineate their functional role in *Candida* infection in intestinal mucosa. Collectively, these findings suggest that Dectin-2 is required for the functional licensing of granulocytes, ensuring effective but balanced effector activity. This interpretation aligns with previous work linking Dectin-2 signalling to neutrophil activation, including integrin upregulation with mediation of anti-fungal defence (22).

The tissue specificity of this defect is striking. While fungal clearance in the kidney was unaffected in Dectin-2 deficient mice, fungal burdens in the intestine failed to resolve, but only in the presence of antigen-specific T cells. This suggests that Dectin-2 has a checkpoint role specifically within mucosal effector sites, where it may tune the responsiveness of neutrophils and eosinophils. Our observations may also help explain discrepancies in prior studies. While Dectin-2 has been shown to be essential for systemic protection in several infection models, other studies reported intact fungal clearance at early time points (5, 8). Our findings extend this view by suggesting that Dectin-2 functions in a tissue- and phase-specific manner, dispensable for the initial innate containment of fungal growth, but required once antigen-specific CD4 T cells are activated and recruited into mucosal tissues. This work also supports the idea that antigen-specific tools are necessary to uncover the precise immune dynamics during fungal infections. Past studies often assessed cytokine production from bulk CD4 T cell populations, but our use of an antigen-specific T cell model allowed us to track antigen-specific T cell behaviour and link it directly to fungal clearance outcomes.

In summary, we demonstrate that Dectin-2 is not required for priming or polarization of CD4 T cells but is critical for regulating Th17 cytokine output influencing granulocyte activation and fungal clearance in the gastrointestinal tract. This work highlights a distinct late-stage effector-phase role for Dectin-2, emphasizing the need to understand PRR function within both the inductive and execution arms of adaptive immunity at tissue sites of fungal dissemination.

## Acknowledgements

We thank the staff of the animal facilities at the University of Exeter for the care and support of our animals.

## Materials and Methods

### Mice

C57BL/6J (WT) and Dectin-2 KO mice (24) were bred and maintained under specific pathogen-free (SPF) conditions at Charles River Laboratories, UK. OT-II transgenic mice stock number 004194 – B6.Cg-Tg(TcraTcrb)425Cbn/J were purchased from Charles River Laboratories and a breeding colony maintained under SPF conditions at the University of Exeter, UK. WT and Dectin-2 KO mice were co-housed for at least 14 days before experiments and used at 8-12 weeks of age. For all studies, age-matched females were randomly assigned to experimental or control groups. All procedures complied with the University of Exeter ethical review process and UK Home Office licence PP9965358.

### *Candida albicans* strain and infection

Mice were challenged intravenously with 1.5 x 10⁵ yeasts of *C. albicans*. For OT-II cell studies, mice were infected 24 h after adoptive transfer with the OVA-expressing *C. albicans* strain Calb-Ag (13). Yeasts were grown in YPD broth at 30 °C for 24 h, washed twice in PBS, counted, and adjusted to the required concentration. At the indicated time points, mice were euthanised and tissues collected. For fungal burden, organs were homogenised in PBS, serially diluted, and plated on YPD agar supplemented with gentamicin (100 µg/ml) and vancomycin (10 µg/ml). Plates were incubated at 37 °C for 24 h before counting CFU.

### OT-II purification for adoptive transfer

For adaptive immune conditions, single-cell suspensions from peripheral lymph nodes (pLNs) of OT-II (CD45.1) mice were prepared in PBS. CD4 T cells were purified by negative selection using the EasySep Mouse CD4+ T Cell Isolation Kit (STEMCELL Technologies), as per manufacturer’s instructions, washed twice, counted using a Vi-Cell (Beckman Coulter), and adjusted to 10 x 10⁶ cells/ml. A total of 1 x 10⁶ cells in 100 µl PBS was injected intravenously (lateral tail vein) into mice. See Figure 1A and C for infection and adoptive T cell transfer schematics.

### Tissue processing and fungal burden

Kidneys and intestines were harvested on days 3 and 6 post-infection. Tissues were homogenised in PBS and plated for colony-forming units (CFUs).

### Flow cytometry

Mesenteric lymph nodes (mLNs), pLNs and spleens were processed through 70 µm filters into RPMI-1640 GlutaMAX. Peripheral blood was collected into EDTA-coated tubes. Lamina propria cell were isolated using the lamina propria dissociation kit (Miltenyi Biotec) according to manufacturer’s instructions. Red blood cells were lysed using BD PharmLyse (BD Biosciences), followed by washing (400 x g, 5 min) and resuspension in RPMI-1640 GlutaMAX with 10 % FCS and 1 % penicillin/streptomycin (complete RPMI) for downstream assays. Cells were stained with antibodies against CD4, CD45.1, CD69, CD63, CD44, Foxp3, CD11b, CD11c, Ly6G, Ly6C, Siglec-F, CD18, CX3CR1, CD80, CD86, CD40, MHC-II (CD4-BUV563/BV510 (RM4-5, RM4-4; BD Biosciences/BioLegend), CD69-BV510 (H1.2F3, BD Biosciences), CD63-PE (NVG-2, BioLegend) CD44-FITC/BV510 (IM7, BD Biosciences/BioLegend), FoxP3-AF488/AF647 (MF23, BD Biosciences), CD11b-BUV395 (M1/70, BD Biosciences), CD11c-BV711 (HL3, BD Biosciences), Ly6G-eFluor450/SB550 (1A8, eBioscience/BioLegend), Ly6C-BV570 (HK1.4, BioLegend), Siglec-F-SB436 (1RNM44N, eBioscience), CD18-APC (H155-78, BioLegend), CX3CR1-BV785/PE (SA011F11, BioLegend), CD80-FITC/BV650 (16-10A1, BioLegend), CD86-AF700 (GL1, eBioscience), CD40-BV605 (3/23, BioLegend), MHC-II-BUV496 (2G9, BD Biosciences). Data were analysed using FlowJo software.

### Cytokine Assays

Intestinal homogenates prepared on day 6 post infection were analysed using multiplex Luminex assays for Cytokines (IL-1β, IL-2, IL-4, IL-5, IL-6, IL-17A, IFN-g, GM-CSF) were quantified using Bio-Plex Pro Mouse Cytokine kits (Bio-Rad) as per manufacturer’s instructions. Plates were read on a MAGPIX system (Luminex), and concentrations were calculated in a Bio-Plex Manager software.

### Statistical Analysis

Statistical comparisons were made using unpaired Student’s *t*-test. p < 0.05 is (*) considered significant, ns = not significant.

---

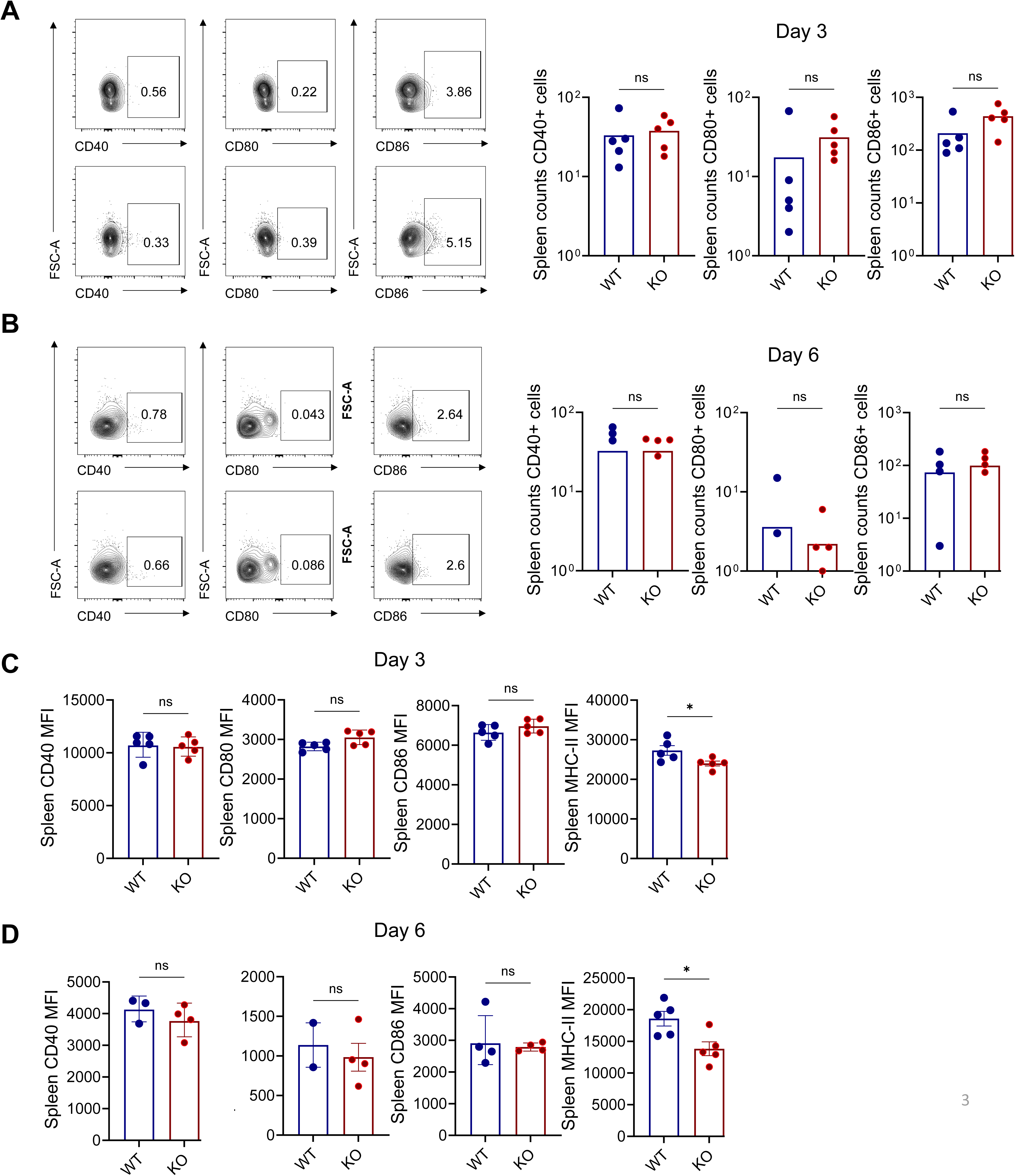
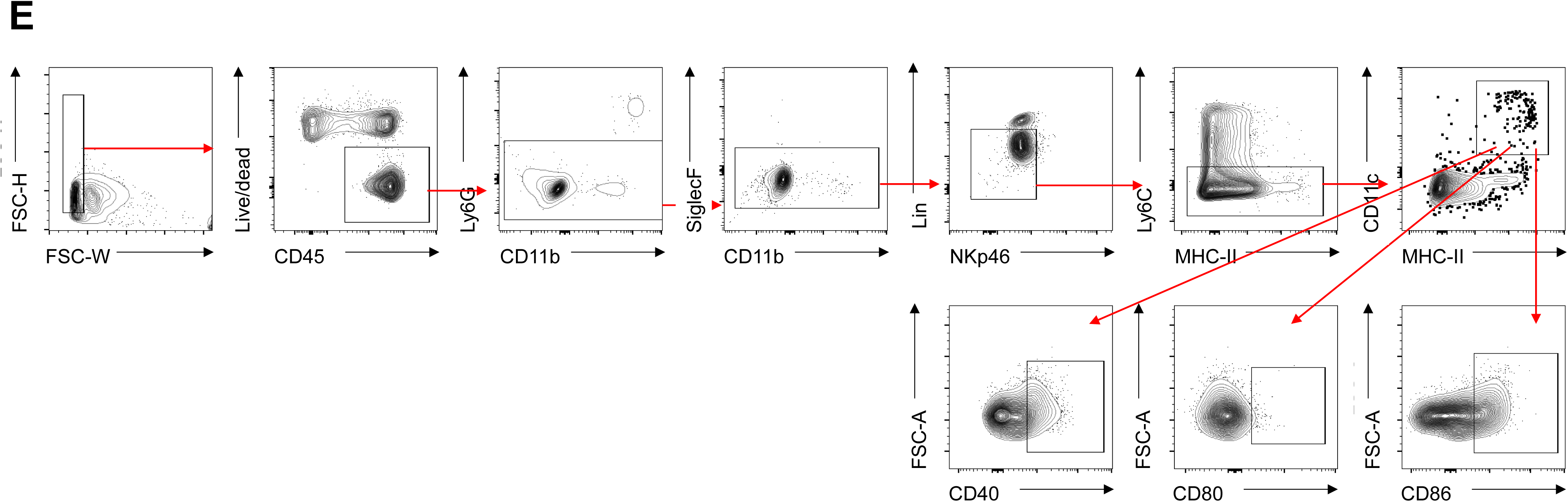

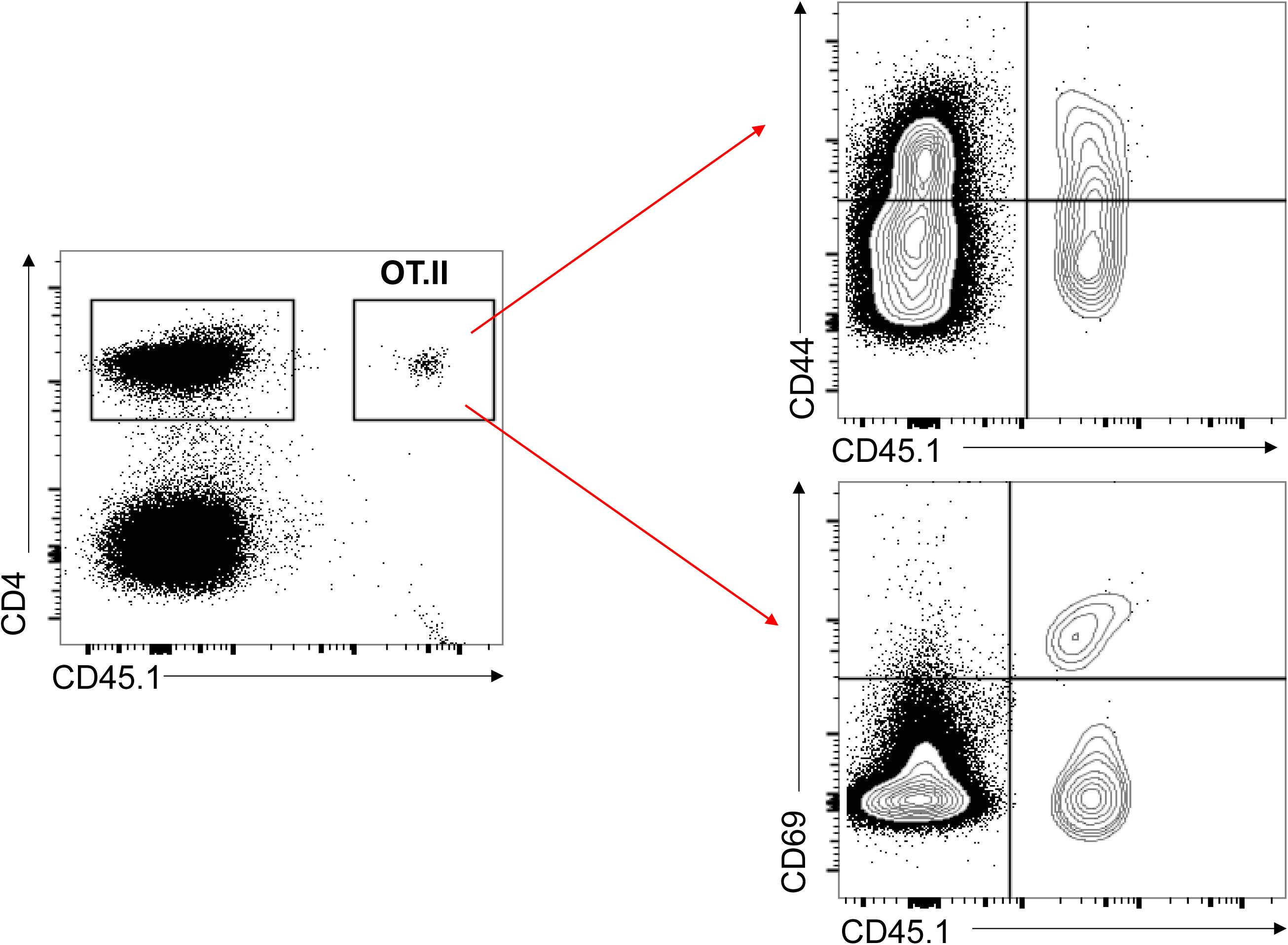

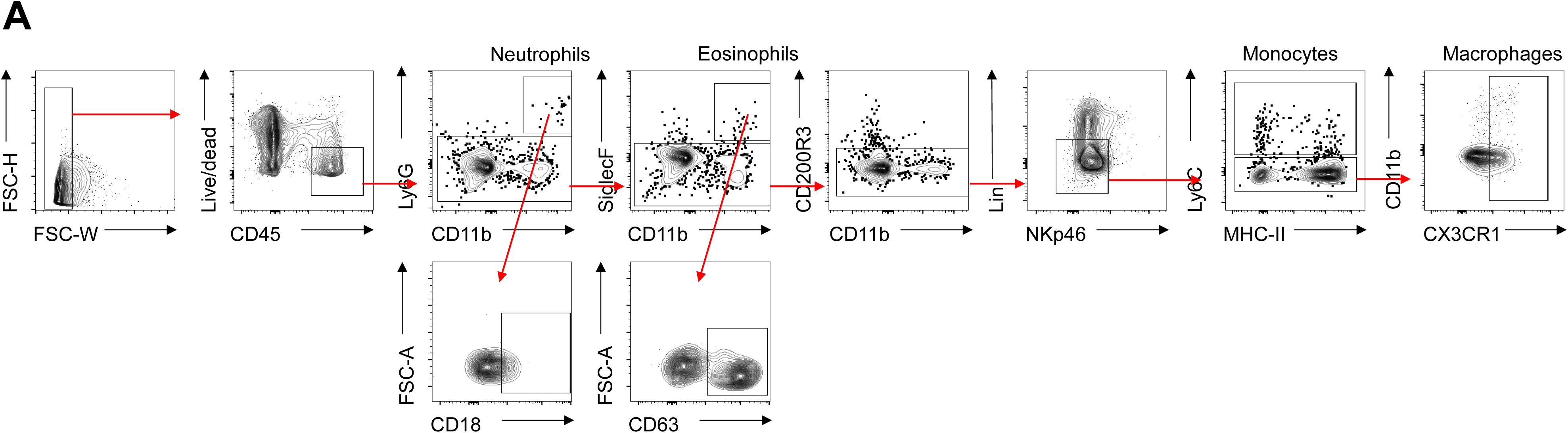

